# Predicting small-molecule inhibition of protein complexes

**DOI:** 10.1101/2024.08.23.609286

**Authors:** Adiba Yaseen, Soumyadip Roy, Naeem Akhter, Asa Ben-Hur, Fayyaz Minhas

## Abstract

**Motivation:** Protein-Protein Interactions (PPIs) are crucial in biological processes and disease mechanisms, underscoring the importance of discovering PPI inhibitors in drug development. Machine learning can expedite this discovery process. Although machine learning techniques for predicting general compound inhibition are available, we are not aware of any that accurately forecast the inhibitory effect of a compound on a *specific* protein complex, utilizing inputs from both the compound and the protein complex.

**Methods:** We present the first *targeted* machine learning based predictor of small molecule based inhibition of protein complexes. Our proposed graph neural network integrates the structure of a protein complex, its protein-protein binding site or interface features and a compound’s SMILES representation to predict the potential of the given compound to inhibit the interaction between proteins in the given complex in a targeted manner.

**Results:** Validated on the 2p2i-DB-v2 database, encompassing 714 inhibitors across 23 complexes with over 12,000 instances, our model achieves superior predictive accuracy (cross-validation AUC-ROC of 0.86), outperforming established kernel methods and pre-trained neural networks. We further tested the predictive performance of our model on two independent external datasets – one collected from recent publications and another consisting of putative inhibitors of the SARS-CoV-2-Spike and Human-ACE2 protein complex with AUC-ROCs of 0.82 and 0.78, respectively. Our targeted predictor introduces a novel approach for PPI inhibitor discovery, laying foundational work for future advancements in addressing this complex and previously unexplored prediction challenge.

**Availability:** Code/supplementary material available: https://github.com/adibayaseen/PPI-Inhibitors

## 1. Introduction

Proteins are involved in precise and targeted interactions with other protein which gives them a diverse set of functions. PPIs play an indispensable role in cellular processes and they are also essential to the mechanisms of numerous cellular functions. However, an undesired interaction can disrupt the function of a target protein. Such pathological interactions can cause diseases like cancer, neurological disorders, and cardiovascular diseases (Kuenemann et al. 2016; Cunningham, Qvit, and Mochly-Rosen 2017; Safari-Alighiarloo et al. 2014; Guo, Wisniewski, and Ji 2014). For example, the B-cell lymphoma/leukemia-2 (Bcl-2) protein is an important controller for planned cell death or apoptosis (Wei et al. 2022). The irregular expression of Bcl-2 is responsible for the development of cancer along with neurodegenerative disorders (Qian et al. 2022). When Bcl-2-associated X protein (Bax), a non-apoptotic protein, interacts with Bcl-2, the formation of the resulting complex (PDB:2xa0) disrupts the normal function of Bcl-2. As a result, uncontrollable cell growth occurs which results in cancer. This characteristic of protein-protein interactions (PPIs) makes them an attractive target for development of novel therapeutic interventions. All such PPIs are druggable targets for small molecule inhibitors (see Fig 1). There are a large number of PPIs in the human proteome but only 2% of them are targeted for drug development indicating an explored frontier for drug development (Gonzalez and Kann 2012) (G. Zhang, Andersen, and Gerona-Navarro 2019).

**Figure 1.**
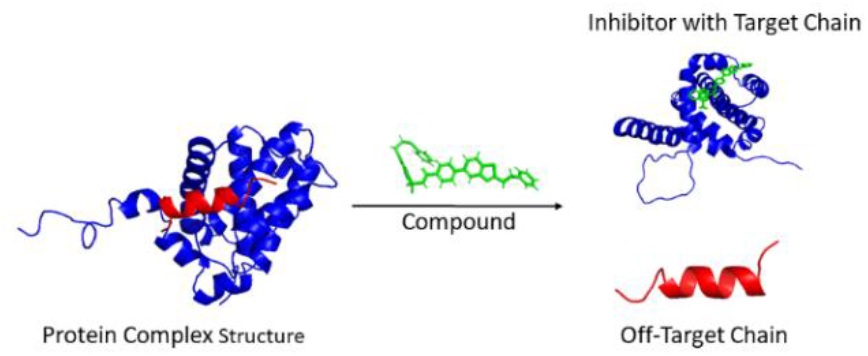
Inhibition of a protein complex by a small molecule (compound): A protein complex consists of two chains (the target chain shown in blue and the off-target chain in red). A given compound can act as an inhibitor for this complex if it can bind to a target chain and disassociate the off-target chain.

Recently, the idea of targeting hot spots in protein interfaces with light weight compounds has emerged (Bogan and Thorn 1998). However, experimental studies for finding inhibitors of PPIs are expensive and laborious so there is room for computational methods to assist the discovery of such compounds.

Docking and molecular dynamics (MD) simulations are two widely used computational methods for identifying top compound hits against a single target protein from large compound databases (Sable and Jois, 2015). However, accurately predicting inhibitors for novel proteins remains challenging due to the poor correlation between top hits and actual inhibitors (Pantsar and Poso, 2018). Additionally, even if docking is successful, it only provides predictions for a single target protein, necessitating the screening of large databases for each new protein with a significant computational overhead. To the best of our knowledge, there is no existing generalized docking method specifically for identifying inhibitors of given protein complexes.

Existing machine learning approaches for this domain are also non-targeted in nature, i.e., they do not predict inhibition of a specific protei complex by a compound. There are a few machines learning methods for PPI inhibitor prediction based on the hypothesis of targeting interface hotspots with light weight compounds. PPIMpred (Jana et al. 2017) and SMMPPI (Gupta and Mohanty 2021) use Support Vector Machines (SVMs) and pdCSM-PPI (Rodrigues, Pires, and Ascher 2021) uses Graph-based signature of encoding of Compounds for this purpose. SMMPPI is the latest method for inhibitor prediction of PPIs which employs a two-stage network, where the initial stage predicts the probability of being an inhibitor against 11 families. However, it is important to note that this model does not utilize protein features and thus cannot act as a protein-complex specific predictor of inhibition.

To the best of our knowledge, all existing machine learning based protein inhibitor prediction methods are non-targeted in nature, i.e., they are unable to predict if a certain small molecule can act as an inhibitor against a specific protein complex or not. This is key question behind this work, i.e., ***predict if a compound can act as an inhibitor of a particular complex*** as shown in Figure 1. There are several additional challenges in accurate inhibitor prediction resulting from the lack of large experimentally verified positive datasets, non-existence of true negative examples, limited availability of protein three-dimensional structures of target proteins, the need for careful experiment design to avoid over-estimation of predictive performance due to the pairwise nature of the underlying prediction problem (Yaseen et al. 2022). In this work, we have aimed to address these challenges.

The **key contributions** of this work are as follows:

1. We emphasize the significance of protein complex-specific or targeted prediction of inhibitors, which constitutes the main research question. We have developed the first complex-specific predictor using a Graph Neural Network (GNN) pipeline that can generate prediction for novel protein complexes.
2. In the absence of experimentally verified negative examples, we used a combination of three different strategies to produce a set of hard and more realistic negative examples to improve the training of the model and its generalization to unseen test examples.
3. To thoroughly assess the performance of our approach, we conducted a rigorous validation, comparing it to a baseline kernel method and protein embeddings from a pre-trained GNN. Additionally, we assessed the effectiveness of our model using independent external datasets.

## 2. Methods

Figure-2 shows the high-level overview of the proposed graph neural network approach which takes a protein complex and a compound as input to predict the inhibition of the protein complex by the compound. Thus, an input example or instance to the model consist of a pair of a protein complex and a compound. Features are extracted from protein chains in the protein complex, their protein-protein binding interface and the chemical structure of the compound. In order to train and validate the machine learning model, we obtain a set of ‘positive’ examples, i.e., examples in which a compound is known to inhibit a complex from 2P2I. As there is currently no gold-standard database of ‘negative’ examples for which it is known that the protein complex is not inhibited by the protein complex, we devised a strategy to generated negative examples that are reflective of the real-world use case for the model. Below, we present details of our datasets, experimental design, and machine learning methods for inhibitor prediction of PPIs.

### 2.1 Datasets

We details of positive and negative examples used in training and validating the model as well as the external test used in independent model assessment are given below.

#### 2.1.1 Positive examples

2P2I v2 (Basse et al. 2016) is the only publicly available database that has structural information on protein complexes and their inhibitors, which is essential for targeted inhibitor prediction. 2P2I contains 32 protein complexes with 822 experimentally verified and manually curated examples of protein complexes and associated ligand; overall the database contains interactions for 733 unique small molecule ligand compounds. Each example consists of a small molecule inhibitor and protein complex such that the inhibitor binds a chain (called the target chain) of a complex and causes it to disassociate from the complex (see Fig 1). Upon analyzing the dataset, we discovered that seven complexes had only predicted structures. To ensure the accuracy of our results, we removed these examples from the positive set, resulting in 722 positive examples. From the remaining 25 complexes, we also removed complexes that have only one inhibitor for robust performance assessment. Our final dataset has 714 examples against 22 protein complexes with 608 unique inhibitors. Details of these protein complexes in terms of their PDB complex identifier, constituent chains and inhibitors as well as associated negative examples are given in Supplementary data Table-1.

**Table 1.**
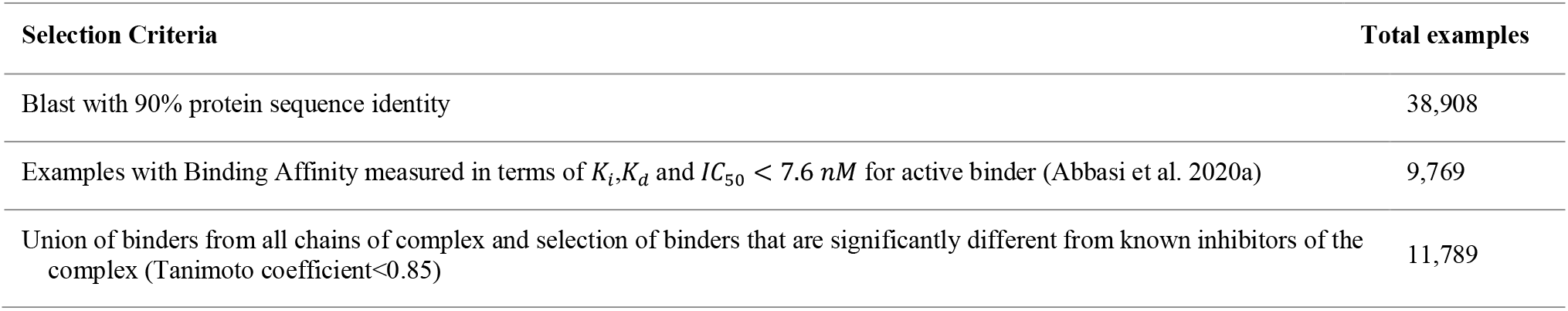
Selection criteria applied to Binding DB for generating the negative dataset.

| Selection Criteria | Total examples |
| --- | --- |
| Blast with 90% protein sequence identity | 38,908 |
| Examples with Binding Affinity measured in terms of $K_i, K_d$ and $IC_{50} < 7.6 \text{ nM}$ for active binder (Abbasi et al. 2020a) | 9,769 |
| Union of binders from all chains of complex and selection of binders that are significantly different from known inhibitors of the complex (Tanimoto coefficient $< 0.85$) | 11,789 |

#### 2.1.2 Generation of negative examples

In the absence of experimentally verified negative examples, i.e., compounds and complex pairs for which it is known that the compound does not inhibit the complex, we resorted to generation of synthetic negative examples. For this purpose, we used a combination of three different strategies to produce a large and “hard” set of negative examples to improve the training of the model and its generalization to unseen test examples as discussed below. These three strategies resulted in a total of 10, 413 negative examples. The first two strategies generated negative examples by pairing protein complexes and compounds in a random manner motivated by the fact that the probability of a randomly selected compound to be an inhibitor for a randomly selected protein complex in the protein universe is quite small. As the number of negative examples is expected to be quite large in comparison to the number of positive examples in the protein and ligand space, we generated 10 times more negative examples as positive examples. However, such negative pairing does not account for the fact that not all compounds that do bind a protein can act inhibitors of complexes involving that protein. To account for this, we generated negative examples consisting of binders of proteins in a protein complex that are, with high probability, not inhibitors of complexes involving these proteins. Each of these three strategies is discussed below:

##### Complexes from 2P2I paired with randomly selected small molecules from 2P2I and SuperDRUG2

We generated negative examples by randomly pairing a complex from 2P2I with other small molecules from 2P2I and SuperDRUG2 (version 2) (Siramshetty et al. 2018) with a total of 3,633 unique small molecules such that the selected compound is not a known inhibitor of that complex. This resulted in a total of 857 examples. (Give total number of examples) negative examples.

##### Compounds from 2P2I paired with complexes from DBD 5

We generated additional negative examples by randomly pairing compounds from 2P2I with 282 complexes from the DBD benchmark database (version 5.5) (Vreven et al. 2015, 2) for which bound three-dimensional structures are available. This resulted in a total of 1714 examples.

##### Generation of binders that are not inhibitors

Using the two aforementioned strategies, random pairing can produce compound and protein complex pairs where the compound might not be a binder of any protein chain in the complex. Relying solely on negative examples from these strategies, along with positive examples of inhibitory compounds, could lead to a biased predictive model. Such a model might struggle to distinguish between compounds that inhibit a protein complex and those that bind to the protein chains without leading to inhibition. It is crucial to note that while all inhibitors bind to the protein complex, not all binders are inhibitors. To mitigate this issue, we incorporate additional ‘hard’ negative examples. These examples consist of compounds known to bind at least one chain in the protein complex but are not recognized inhibitors. This is achieved using the Binding-DB database (Gilson et al. 2016). For each chain in the complex of a positive example, we performed a BLASTp search in Binding-DB at a threshold of >90% sequence identity to identify binding ligands for that chain which have binding affinity measured in terms of *K*_*i*_,*K*_*d*_ and *IC*_50_ < 7.6 *nM* (Abbasi et al. 2020b). In order to exclude possible inhibitors, we only keep those binding ligands that have a Tanimoto coefficient of < 0.85 with any known inhibitors of that complex. The set of all such ligands is then paired with the complex to produce negative examples. Table-1 shows the number of ligands which are obtained after applying the various filtering steps to yield the final set of negative examples.

#### 2.1.3 Independent Test sets

##### Test set extracted from recent publications

In order to test the effectiveness of our proposed method, we collected an independent external dataset consisting of newly discovered inhibitors reported in the literature that have distinct structures compared to those in our cross-validation dataset. We collected a total of 28 inhibitors along with the PPI complex and target chain. Supplementary data Table-3 provides further information regarding this external dataset. Negative examples were generated by pairing a test complex with compounds using the aforementioned negative example generation strategies. The set of positive and negative examples for this dataset are available in Supplementary data.

##### SARS-CoV-2 inhibitors

For further performance analysis, we also collected a set of 25 inhibitors of the RBD-hACE2 PPI that were experimentally identified (Hanson et al. 2020). Negative examples were generated using the aforementioned negative example generation strategies. The set of positive and negative examples for this dataset are available in Supplementary data Table-4.

### 2.2 Feature Extraction

Each prediction of targeted protein complex inhibition involves a ligand or compound and a protein complex, comprising multiple protein chains. We derive diverse features from ligands, protein complexes, their constituent proteins, and their protein-protein binding sites. These features serve as inputs to a graph neural network model, which generates embeddings for protein complex structures. These embeddings are integrated with interface, protein sequence, and ligand features to produce predictions (refer to Fig 2). This section outlines the types of features extracted from the components of each example.

**Figure 2.**
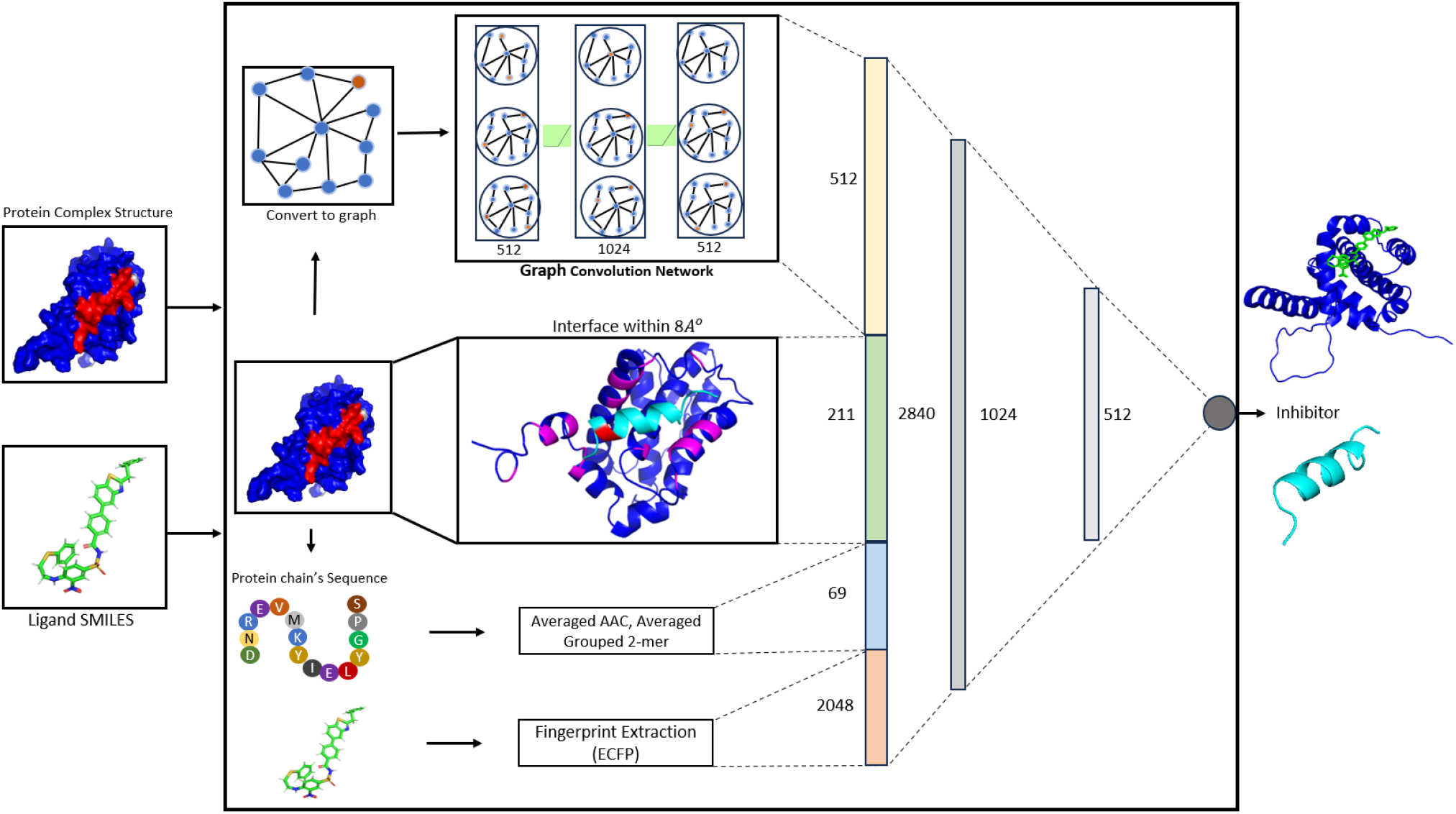
The proposed Graph Neural Network-based model. The model takes the 3D structure of the protein complex consisting of two chains, with the target chain shown in red which is expected to bind the compound and blue being the non-target chain. The model also takes and the SMILES representation of the ligand as input to produce an inhibition potential score for this example. The model builds a feature representation for the protein complex using a GNN which is coupled with amino acid composition features of the overall protein sequence as well as features derived from residues in the interface of the protein complex (defined by amino acids on the two chains that are within 8Å). The ligand (compound) is represented using its DeMorgan/ Extended-Connectivity Fingerprints (ECFP) Fingerprint representation. All features are concatenated to produce a 2,840 dimensional feature representation which is passed to a multilayer perceptron (MLP) to generate the final output after end-to-end training. The generated score reflects the ability of the ligand to disassociate/inhibit the protein complex

#### 2.2.1 Ligand Features

For modeling the compounds, we extract features from their SMILES string representation in the form of the Extended-Connectivity Fingerprint (ECFP), also known as Morgan Features (M. Veselinovic et al. 2015) with the RDKit framework (K. Huang et al. 2020). This method is commonly used for encoding compound information in cheminformatics and drug discovery. It captures the structural information of molecules by encoding the presence or absence of substructural fragments within a given radius of each atom in the molecule. With a radius of 2 bonds, ECFP considers the immediate neighbors of each atom and the atoms directly connected to them, leading to a feature representation with 2,048 dimensions.

#### 2.2.2 Protein Sequence Features

For the purpose of capturing amino acid-specific binding characteristics of constituent proteins chains, we have utilized the amino acid composition (AAC) of a protein and grouped k-mer composition features. AAC is a 20-dimensional vector containing the frequency of occurrence of amino acids in a protein sequence (K. Huang et al. 2020).

We also captured the physiochemical similarity between amino acids through grouped k-mer representation of proteins as features of each protein. In this method, an amino acid is assigned to one of seven amino acid groups based on its physicochemical characteristics (as shown in supplementary Table-2) (Hashemifar et al. 2018) and then the counts of grouped k-mers in a protein are used as features. For *k* = 2, this results in 7^2^ = 49 features of a protein chain. As a protein complex consists of multiple protein chains, the protein sequence features are averaged across protein chains in the same complex.

#### 2.2.3 Interface Features

PPIs are a consequence of non-covalent interactions between interface residues of a target chain with off-target chains in a complex. Consequently, features extracted from the protein-protein interface can be very useful in predicting PPI inhibitors. We calculated the number of unique pairs of residues at the interface of a protein complex based on its 3D structure. More specifically, we computed a 211-dimensional feature vector where each element represents the frequency of occurrence of a specific amino acid pair in the interface (including an indictor amino acid label for non-standard or unspecified amino acids). The interface residues are identified as residues found in both protein chains that are within a distance of 8 Angstroms from each other, as depicted in Figure 2.

### 2.3 A GNN-based inhibitor prediction model

We have developed a graph neural network-based model that takes the three-dimensional structure of a protein complex as input and produces an embedding for it which is then combined with other protein interface, sequence-based features and the compound features discussed above (see Fig 2). The combined embedding is then passed through two fully connected layers to produce the final prediction score. The entire model is trained in an end-to-end manner. By integrating different types of features, including sequence and structure, our model exploits their synergistic nature, enabling a more comprehensive representation of protein interactions.

#### 2.3.1 A Heterogeneous Graph representation of protein complexes

The protein complex structure is converted into a binary contact map and fed to a graph neural network (GNN) for inhibitor prediction of PPIs. The GNN is heterogeneous in that it incorporates both atomic contacts as well as residue level features. Let *G* (*V, E*) denote the graph representing the complex, where each node (*𝒱* ∈ *V*) is the representation of an atom in the protein and edges in the set *E* represent atomic contacts. Two atoms are considered as connected if their distance is less than 6 Angstroms as shown in Figure 3.

**Figure 3.**
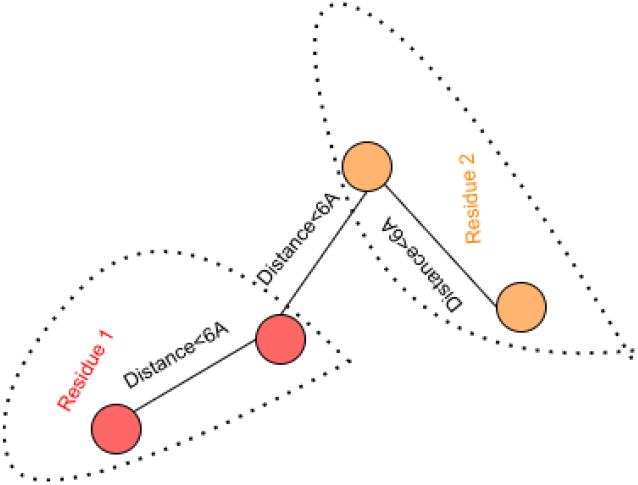
Information of same and different neighbors within a protein structure used in the graph neural network

#### 2.3.2 Node features of atoms and residues

Each node has a vector of node features in the graph of a protein complex which are derived from the protein complex structures and its sequence. We represent each atom and residue using their respective one hot encoding (OHE). An atom can belong to one of the following categories: {*C, CA, CB, CGG, CH*^2^, *N, NH*^2^, *OG, OH, O*^1^, *O*^2^, *SE*} (each corresponding to a distinct atom type or group) resulting in a 12 dimensional vector representation 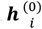 of each atom indexed by subscript *j*. If an atom does not belong to one of the above atom types, we set: 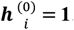.

Each node is also associated with the one-hot encoding of the residue that it belongs to. This residue level one-hot encoding is denoted by ***r***_*i*_ which encodes each residue into 21 different categories of amino acids (the last one to represent unknown amino acids).

#### 2.3.3 Graph neural network layers

We have used a 3-layer GNN which takes the atomic 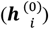 and residue (***r***_*i*_) level embeddings of the protein structure to build higher-order embeddings of individual atoms in the protein and consequently the entire protein. For this purpose, the first layer of the GNN builds a representation 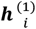 of each atom in the protein using the OHE vector representation of the atom itself, the residue to which that atom belongs and of 10 nearest neighboring atoms within the same residue as well as 10 nearest neighboring from other residues (see Fig 3). Each subsequent layer of the GNN then takes the adjacency list and node embeddings from the previous layer 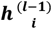 as input and outputs the node-level the embeddings for next layer 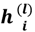 as described in the following equations:

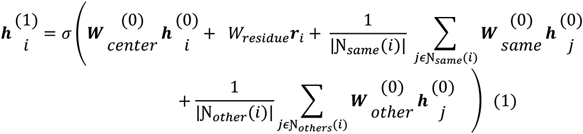

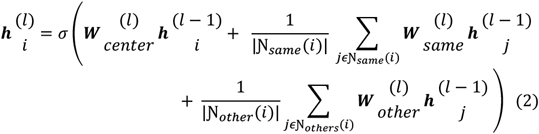

Here:

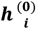 : atomic one-hot-encoding for atom *i*

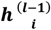 : embedding of atoms from the (*l* − 1)*th* layer

***r***_***i***_: residue-level one-hot-encoding for residue containing atom *i*

Ɲ_***same***_(***i***): Set of neighboring atoms of atom *i* that are within the same residue

Ɲ_***other***_(***i***): the set of neighboring atoms of atom *i* that are from different residues

σ: the activation functions (ReLU).

The weight matrices (denoted by ***W***) are learnable parameters of the model which are used to produce atomic embeddings for each layer of 512,1024 and 512 dimensions for the three layers. The resulting embeddings are then aggregated to generate the protein level embedding.

#### 2.3.6 Overall network structure and training

The GNN layers produce a 512-dimensional vector representation of a protein which, in conjunction with the hand-crafted sequence and interface-based features are passed to a fully connected (FC) layer with a hyperbolic tangent (tanh) activation in the first two layers and ReLU in the third layer as shown in Figure 2. The dimension of all concatenated features representing a single example becomes 2840 which are passed through a two hidden layer multi-layered perceptron with 512 and 100 neurons to produce a single prediction score representing the inhibition potential of the example. We use the Binary Cross Entropy loss in combination with a learning rate of 0.0001 and Adam optimizer. Due to the significant class imbalance in the dataset, we have utilized a weighted averaging strategy in model training in which errors over positive examples are weighted more than those over negative examples with the weighting determined by the positive to negative class ratio for each complex.

### 2.4 A Heterogeneous Kernel-based baseline model

As a baseline, we have also developed a simple kernel-based method for inhibitor prediction of PPIs based on our previous paper for compound protein interaction prediction (Yaseen et al. 2022). As each classification example in this problem comprises a protein complex and compound, we first construct non-linear radial basis function (RBF) similarity kernel representations of protein complex features and compounds separately based on their respective features which are then combined to form a heterogeneous feature space kernel for classification as shown in Figure 4 (for further details, see (Yaseen et al. 2022)). This joint kernel representation measures the extent of similarity between two examples with each example being a protein complex-and compound pair. Note that the joint kernel is a product of the protein and compound kernels which gives rise to an abstract joint feature space indirectly resembles to the tensor-product of the protein and compound feature spaces. It is also significant to understand that two samples will have a high kernel score if the corresponding complex and inhibitor in the two examples are similar. The resulting kernel is then passed to a support vector machine for classification with a custom kernel.

**Figure 4.**
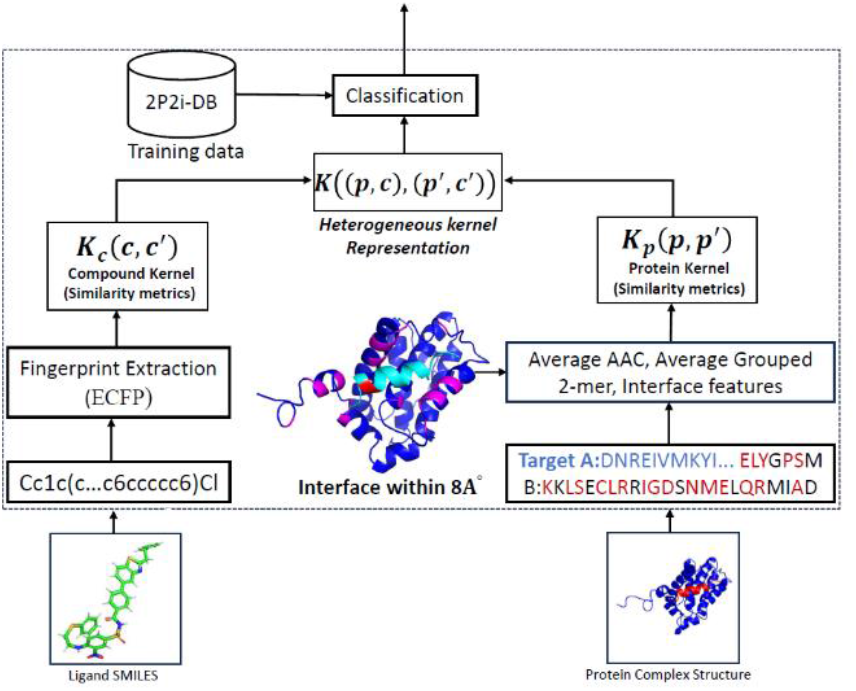
Concept diagram of Heterogenous Kernel-based Inhibitor of Protein-Protein Interaction (Kernel-IPPI) Prediction. 3D-structure of Protein consisting of 2 chains (red and blue) and SMILES representation of a ligand are given as input. Interface features consisting of amino acid composition of residues on the two chains within 8 Angstroms as well as the overall amino acid composition and grouped k-mer (k = 2) features are extracted from the protein complex. These features are used for constructing a protein-level kernel ***K***_***p***_(***p, p***^′^). The Morgan Fingerprint is computed from SMILES representation of a compound to construct the compound kernel ***K***_***c***_(***c, c***^′^). These kernels are concatenated into a kernel vector ***K***((***p, c***), (***p***^′^, ***c***^′^)) for prediction via an SVM.

### 2.5 GearNet-Edge Feature Embeddings

In order to test the effectiveness of the proposed approach in the context of existing graph based pipelines available for different protein function prediction tasks, we used the embeddings of protein complexes obtained from the Geometry-Aware Relational Graph Neural Network (GearNet) model (Z. Zhang et al. 2022) with a multi-layered perceptron for classification. GearNet-Edge uses a residue graph with multiple relations along with mult-view contrastive loss learning aimed to enhance the similarity between distinct sub-structures within the same protein 3D structure while minimizing similarities between different protein complexes. The geometric encoder utilizes edge messaging passing for macromolecular representation learning. We have used the publicly accessible codes of GearNet -Edge for performing an experiment and comparison with our GNN-based method.

## 3 Results and Discussion

### 3.1 Leave One Complex Out Cross Evaluation

To obtain a realistic evaluation of performance, we have used Leave-One-Complex-Out Cross-Validation (LOCO) analysis. This approach involves excluding all examples associated with a complex from the training set and utilizing them as a test set to evaluate model performance after training the model on all other examples from other complexes. As discussed in the Dataset section, we have used experimentally verified positive examples from the 2P2I database and negative examples generated using random pairing of both compounds and protein complexes along with binders from Binding DB. Table-2 shows the results for each held-out complex from all methods. The resulting ROC and PR curves are shown in Figure 5. This comparison demonstrates that the proposed GNN-based method shows improved performance in comparison to baseline methods in terms of both average AUROC (0.863) and AUC-PR (0.39).

**Table 2.**
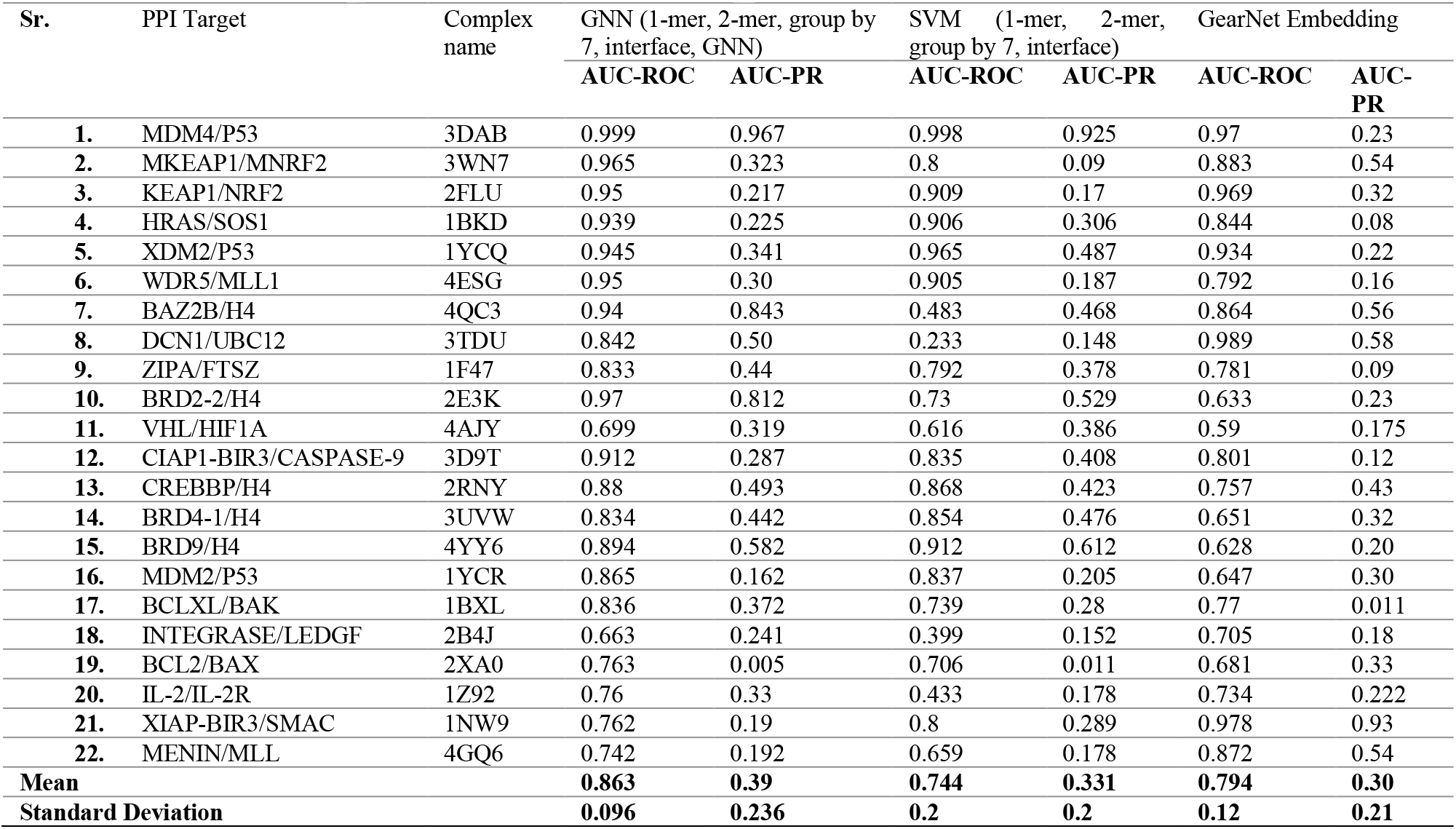
Results of leave one complex out validation across different complexes.

| Sr. | PPI Target | Complex name | GNN (1-mer, 2-mer, group by 7, interface, GNN) |  | SVM (1-mer, 2-mer, group by 7, interface) |  | GearNet Embedding |  |
| --- | --- | --- | --- | --- | --- | --- | --- | --- |
|  |  |  | AUC-ROC | AUC-PR | AUC-ROC | AUC-PR | AUC-ROC | AUC-PR |
| 1. | MDM4/P53 | 3DAB | 0.999 | 0.967 | 0.998 | 0.925 | 0.97 | 0.23 |
| 2. | MKEAP1/MNRF2 | 3WN7 | 0.965 | 0.323 | 0.8 | 0.09 | 0.883 | 0.54 |
| 3. | KEAP1/NRF2 | 2FLU | 0.95 | 0.217 | 0.909 | 0.17 | 0.969 | 0.32 |
| 4. | HRAS/SOS1 | 1BKD | 0.939 | 0.225 | 0.906 | 0.306 | 0.844 | 0.08 |
| 5. | XDM2/P53 | 1YCQ | 0.945 | 0.341 | 0.965 | 0.487 | 0.934 | 0.22 |
| 6. | WDR5/MLL1 | 4ESG | 0.95 | 0.30 | 0.905 | 0.187 | 0.792 | 0.16 |
| 7. | BAZ2B/H4 | 4QC3 | 0.94 | 0.843 | 0.483 | 0.468 | 0.864 | 0.56 |
| 8. | DCN1/UBC12 | 3TDU | 0.842 | 0.50 | 0.233 | 0.148 | 0.989 | 0.58 |
| 9. | ZIPA/FTSZ | 1F47 | 0.833 | 0.44 | 0.792 | 0.378 | 0.781 | 0.09 |
| 10. | BRD2-2/H4 | 2E3K | 0.97 | 0.812 | 0.73 | 0.529 | 0.633 | 0.23 |
| 11. | VHL/HIF1A | 4AJY | 0.699 | 0.319 | 0.616 | 0.386 | 0.59 | 0.175 |
| 12. | CIAP1-BIR3/CASPASE-9 | 3D9T | 0.912 | 0.287 | 0.835 | 0.408 | 0.801 | 0.12 |
| 13. | CREBBP/H4 | 2RNY | 0.88 | 0.493 | 0.868 | 0.423 | 0.757 | 0.43 |
| 14. | BRD4-1/H4 | 3UVW | 0.834 | 0.442 | 0.854 | 0.476 | 0.651 | 0.32 |
| 15. | BRD9/H4 | 4YY6 | 0.894 | 0.582 | 0.912 | 0.612 | 0.628 | 0.20 |
| 16. | MDM2/P53 | 1YCR | 0.865 | 0.162 | 0.837 | 0.205 | 0.647 | 0.30 |
| 17. | BCLXL/BAK | 1BXL | 0.836 | 0.372 | 0.739 | 0.28 | 0.77 | 0.011 |
| 18. | INTEGRASE/LEDGF | 2B4J | 0.663 | 0.241 | 0.399 | 0.152 | 0.705 | 0.18 |
| 19. | BCL2/BAX | 2XA0 | 0.763 | 0.005 | 0.706 | 0.011 | 0.681 | 0.33 |
| 20. | IL-2/IL-2R | 1Z92 | 0.76 | 0.33 | 0.433 | 0.178 | 0.734 | 0.222 |
| 21. | XIAP-BIR3/SMAC | 1NW9 | 0.762 | 0.19 | 0.8 | 0.289 | 0.978 | 0.93 |
| 22. | MENIN/MLL | 4GQ6 | 0.742 | 0.192 | 0.659 | 0.178 | 0.872 | 0.54 |
| Mean |  |  | **0.863** | **0.39** | **0.744** | **0.331** | **0.794** | **0.30** |
| Standard Deviation |  |  | **0.096** | **0.236** | **0.2** | **0.2** | **0.12** | **0.21** |

**Figure 5.**
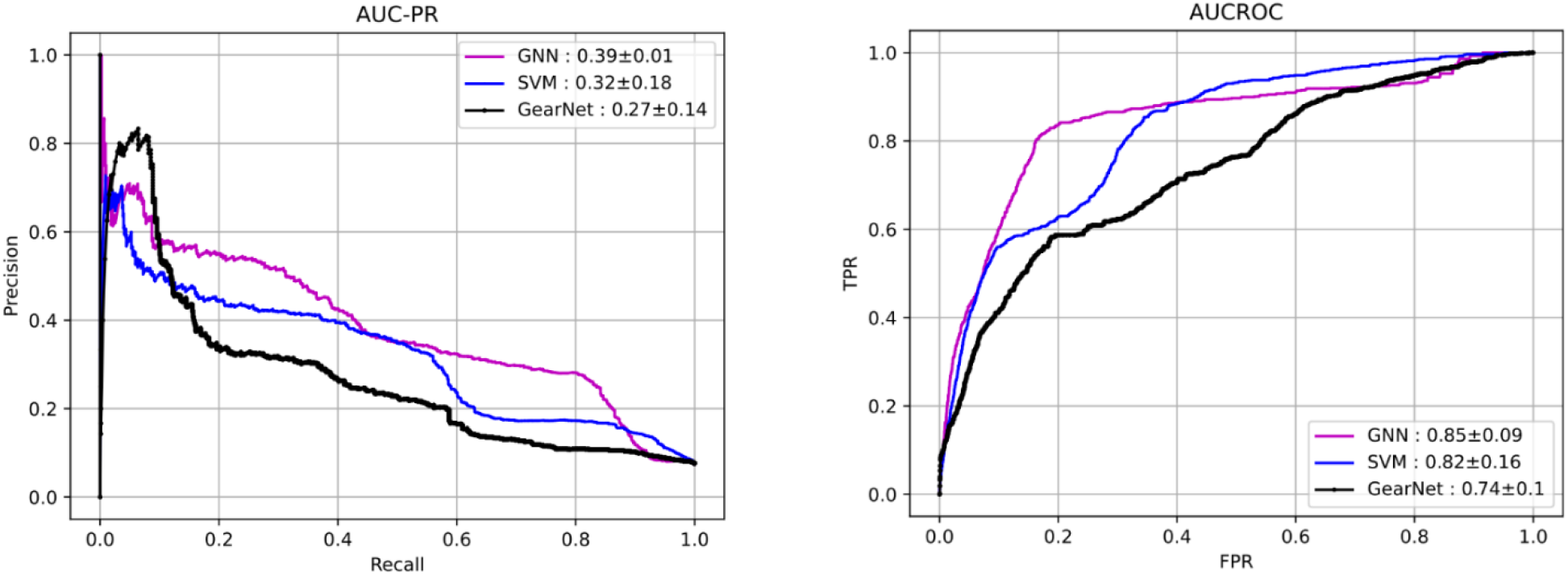
Precision-Recall (PR) (left) and Receiver Operating Characteristic (ROC) (right) curves of leave one complex out validation for our baseline method (SVM), proposed Graph neural network-based method in combination with GearNet based results. Reported are also the average AUCs along with standard deviations of each method

### 3.2 External evaluation

In order to assess the prediction quality for unseen data we collected examples from recent publication which consists of protein complexes with low sequence similarity to our training data. Random negative examples are generated using the FDA-approved drugs in the SuperDRUG2 dataset. We also use DBD5 complexes for pairing inhibitors. All these examples from latest publications are made available to the community as supplementary material. We evaluate the robustness of the GNN-based method for inhibitor prediction of unknown protein target and it achieves 82% AUC-ROC as shown in Figure 5. We also evaluate our model for COVID-9 data protein complexes from the study by Hanson et al. The proposed model achieves an AUC-ROC of 76% for these examples as shown in Figure 6. These results clearly show that the proposed approach can effectively predict targeted inhibitors of protein complexes.

**Figure 6.**
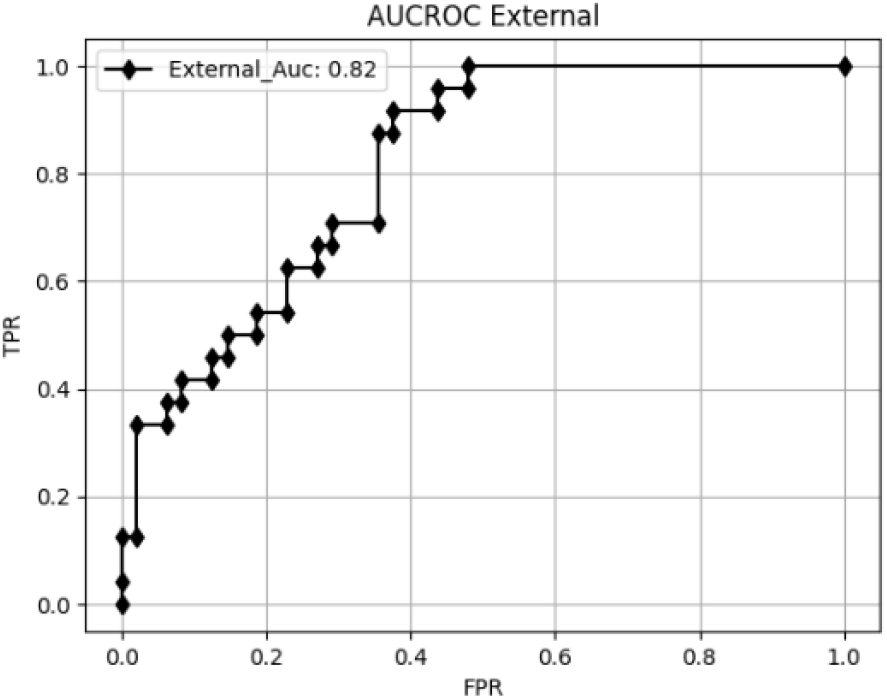
External evaluation for inhibitor prediction for Noval proteins collected from recent publication.

**Figure 7.**
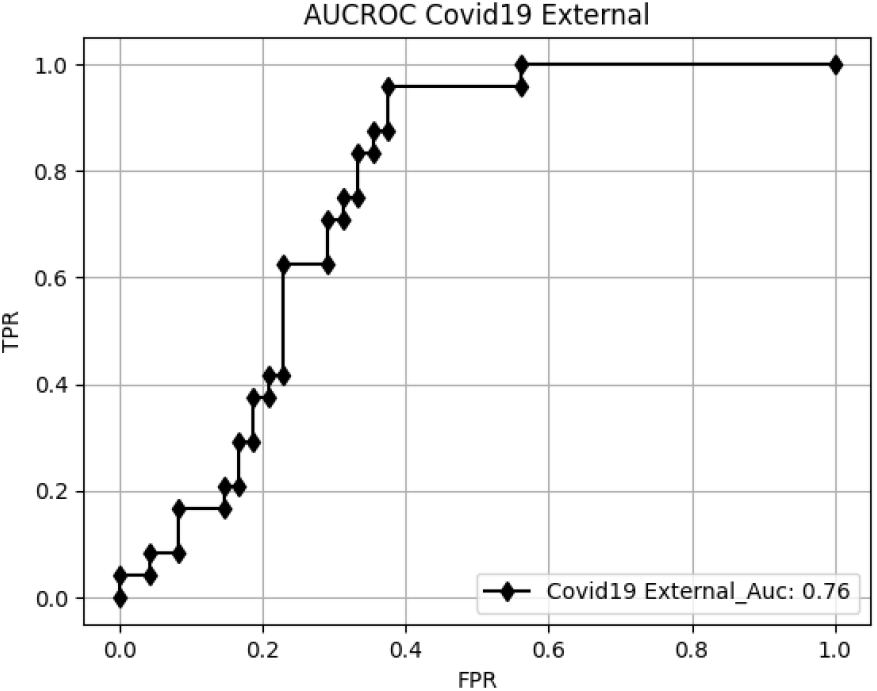
Inhibitor prediction for putative chemical ligands of SARS-CoV-2-Spike and Human-ACE2 proteins

## 4 Conclusions

In this paper, we have aimed to address a critical shortcoming of existing methods for predicting small molecule inhibitors of protein complexes by presenting a GNN-based method that can predict the inhibition potential of a small molecule for a target protein complex. We show that the proposed method offers superior performance compared to other baseline methods in both cross-validation as well as external test sets. This study can help in refining drug development strategies especially for diseases involving protein-protein interactions and paves the work for further development in this previously unexplored prediction problem.

## Acknowledgments

This work is supported by Pakistan HEC NRPU 6085.

