## supplementary for "Predicting small-molecule inhibition of protein complexes"

### Supplementary material of Predicting small-molecule inhibition of protein complexes

| Table 1 Detail of 2P2I Dataset | | | | | | |
|---|---|---|---|---|---|---|
| Sr. No | PPI Target | Complex Name | Positive | Negative | N/P ratio | Names of chains |
| 1 | 'TNFA/TNFA | 1TNF | 1 | - | - | ['A', 'B'] |
| 2 | TNFR1A/TNFB | 1TNR | 1 | - | - | ['B', 'A'] |
| 3 | HPV-E2/E1 | 1TUE | 1 | - | - | ['A', 'B'] |
| 4 | XDM2/P53 | 1YCQ | 11 | 320 | 29.1 | [ 'A', 'B'] |
| 5 | INTEGRASE/LEDGF | 2B4J | 65 | 267 | 4.1 | [ 'A', 'B'] |
| 6 | MDM4/P53 | 3DAB | 5 | 314 | 62.8 | ['A', 'B'] |
| 7 | CREBBP/H4 | 2RNY | 61 | 324 | 5.3 | ['A', 'B'] |
| 8 | BRD4-1/H4 | 3UVW | 201 | 1061 | 5.3 | ['A', 'B'] |
| 9 | MDM2/P53 | 1YCR | 51 | 1220 | 23.9 | ['A', 'B'] |
| 10 | IL-2/IL-2R | 1Z92 | 7 | 30 | 4.3 | ['A', 'B'] |
| 11 | XIAP-BIR3/SMAC | 1NW9 | 13 | 144 | 11.1 | ['A', 'B'] |
| 12 | ZIPA/FTSZ | 1F47 | 4 | 17 | 4.2 | ['B', 'A'] |
| 13 | MENIN/MLL | 4GQ6 | 23 | 173 | 7.5 | ['B', 'A'] |
| 14 | BCLXL/BAK | 1BXL | 12 | 65 | 5.4 | ['B', 'A'] |
| 15 | BCL2/BAX | 2XA0 | 3 | 1616 | 538.7 | ['B', 'A'] |
| 16 | HRAS/SOS1 | 1BKD | 2 | 90 | 45 | ['R', 'S'] |
| 17 | KEAP1/NRF2 | 2FLU | 12 | 508 | 42.3 | ['X', 'P'] |
| 18 | BRD9/H4 | 4YY6 | 22 | 100 | 4.5 | ['Z', 'A'] |
| 19 | BAZ2B/H4 | 4QC3 | 104 | 295 | 2.8 | ['C', 'B', 'A'] |
| 20 | VHL/HIF1A | 4AJY | 90 | 279 | 3.1 | ['C', 'V', 'H', 'B'] |
| 21 | WDR5/MLL1 | 4ESG | 30 | 1160 | 38.7 | ['D', 'C', 'A', 'B'] |
| 22 | CIAP1-BIR3/CASPASE-9 | 3D9T | 28 | 633 | 22.6 | ['C', 'A', 'D', 'B'] |
| 23 | MKEAP1/MNRF2 | 3WN7 | 27 | 1446 | 53.6 | ['L', 'A', 'M', 'B'] |
| 24 | BRD2-2/H4 | 2E3K | 66 | 281 | 4.3 | ['R', 'B', 'A', 'C', 'Q', 'D'] |
| 25 | DCN1/UBC12 | 3TDU | 20 | 70 | 3.5 | ['E', 'F', 'A', 'D', 'B', 'C'] |

| Table 2 Amino acid seven classes partition based on their physicochemical characteristics. | | | | |
|---|---|---|---|---|
| Class | Amino acid | Dipole moment | Volume ($A^{o}3$) | Diffused bond creation |
| 1 | A, V, G | <1.0 | <50 | No |
| 2 | I, F, L, P | <1.0 | >50 | No |
| 3 | M, S, T, Y | (1.0, 2.0) | >50 | No |
| 4 | H, N, Q, W | (2.0, 3.0) | >50 | No |
| 5 | K, Rs | >3.0 | >50 | No |
| 6 | D, E | >3.0(opposite direction) | >50 | No |
| 7 | C | <1.0 | <50 | Yes |

| Table 3 External dataset collect from recent publications | | | |
|---|---|---|---|
| Sr. No | Inhibited Complex | Inhibitor Name | SMILES |
| 1 | 7P58 | 5UI | CC(C)Oc1cccc(c1)c2cccc(n2)n3c(c(cn3)C(=O)O)C(F)(F)F |
| 2 | 7P5E | 5VX | CN(C)C(=O)c1cccc(c1)c2cccc(n2)n3c(c(cn3)C(=O)O)C(F)(F)F |
| 3 | 7P5F | 5QZ | CN(C)C(=O)c1cccc(c1)c2cccc(c2)n3c(c(cn3)C(=O)O)C4CC4 |
| 4 | 7P5I | 5X9 | Cn1cc(nn1)C2CC2c3c(cnn3c4cccc(c4)c5cccc(c5)C(=O)N(C)C)C(=O)O |
| 5 | 7P5K | 5UZ | CCCC1CCCCN1C(=O)c2cccc(c2F)c3cccc(c3)n4c(c(cn4)C(=O)O)C5CC5 |
| 6 | 7P5N | 5RQ | CCCCC1CCCN1C(=O)C2CCCC(C2)c3cccc(c3)n4c(c(cn4)C(=O)O)C5CC5 |
| 7 | 7P5P | 5RB | CCCC1CCCCN1C(=O)c2cccc(c2)c3cccc(c3)n4c(c(cn4)C(=O)O)C5CC5c6cn(nn6)C |
| 8 | 6QMC | J6H | c1cc(ccc1C(CC(=O)O)C2=CC(=O)NC=C2)Cl |
| 9 | 6QMD | J6N | Cn1c2ccc(cc2nn1)C(CC(=O)O)c3ccc(cc3)Cl |
| 10 | 6QME | J6Q | Cc1cc(ccc1Cl)C(CC(=O)O)c2ccc3c(c2)nnn3C |
| 11 | 6QMJ | J6K | Cc1ccc(cc1CN(C)S(=O)(=O)c2ccccc2)C(CC(=O)O)c3cc4c(c(c3)OC)n(nn4)C |
| 12 | 6QMK | J8H | Cc1ccc(cc1CN2CCOc3ccccc3S2(=O)=O)C(CC(=O)O)c4cc5c(c(c4)OC)n(nn5)C |
| 13 | 5FNQ | S0W | c1cc(ccc1CCC(=O)O)Cl |
| 14 | 5FNR | XMS | Cn1c2ccc(cc2nn1)C(CC(=O)O)c3ccc(cc3)Cl |
| 15 | 5FNS | XYY | Cn1c2ccc(cc2nn1)C(CC(=O)O)c3ccc(c(c3)CN(C)S(=O)(=O)C)Cl |
| 16 | 5FNT | 0PL | Cn1c2ccc(cc2nn1)C(CC(=O)O)c3ccc(c(c3)CN(C)S(=O)(=O)c4ccccc4)Cl |
| 17 | 5FNU | L6I | Cc1ccc(cc1CN2CC(Oc3ccccc3S2(=O)=O)C)C(CC(=O)O)c4cc5c(c(c4)OC)n(nn5)C |

| | | | |
|---|---|---|---|
| 18 | 5FZJ | 75K | Cc1nc2c(o1)C=C(OC2=O)C |
| 19 | 5FZN | FB2 | c1ccc(cc1)S(=O)(=O)N |
| 20 | 6V6Z | Q5Y | COc1ccc(cc1)S(=O)(=O)Nc2ccc(c3c2cccc3)N(CC(=O)O)S(=O)(=O)c4ccc(cc4)OC |
| 21 | 7XM5 | 6i | O=S(N(CC(N)=O)C1=C2C(C=CC=C2)=C(N(CC(N)=O)S(=O)(C3=CC=C(NC(CCN4CCOCC4)=O)C=C3)=O)C=C1)(C5=CC=C(OC)C=C5)=O |
| 22 | 7XM3 | 6k | O=S(N(CC(N)=O)C1=C2C(C=CC=C2)=C(N(CC(N)=O)S(=O)(C3=CC=C(NC(CCN4CCN(CC)CC4)=O)C=C3)=O)C=C1)(C5=CC=C(OC)C=C5)=O |
| 23 | 7XM4 | 6e | O=S(N(CC(N)=O)C1=C2C(C=CC=C2)=C(N(CC(N)=O)S(=O)(C3=CC=C(NC(CN4CCN(CC)CC4)=O)C=C3)=O)C=C1)(C5=CC=C(OC)C=C5)=O |
| 24 | 7XM2 | GFD | COc1ccc(cc1)S(=O)(=O)N(CC(=O)N)c2ccc(c3c2cccc3)N(CC(=O)N)S(=O)(=O)c4ccc(cc4)N |

**Table-4 inhibitors of the RBD-hACE2 PPI that were experimentally identified (Hanson et al. 2020)**

| Sr. No | Inhibitor Name | SMILES |
|---|---|---|
| 1 | Ciclopirox | CC=2C=C(C1CCCCC1)N(O)C(=O)C=2.NCCO |
| 2 | CetylpyridiniumBromide | [Br-].CCCCCCCCCCCCCCCC[n+]1ccccc1 |
| 3 | CETYLPYRIDINIUM | CCCCCCCCCCCCCCCC[N+]1=CC=CC=C1 |
| 4 | SodiumdeoxycholateMonohydrate | C[C@H](CCC([O-])=O)[C@H]1CC[C@H]2[C@@H]3CC[C@@H]4C[C@H](O)CC[C@]4(C)[C@H]3C[C@H](O)[C@]12C |
| 5 | Triethylenetetramine | NCCNCCNCCN |
| 6 | Hinokitiol | CC(C)C\1=C\C=C/C(=O)\C(=C1)O |
| 7 | Oxfendazole | O=C(OC)Nc1nc2cc(ccc2n1)S(=O)c3ccccc3 |
| 8 | Prulifloxacin | CC=5OC(=O)OC=5CN1CCN(CC1)c2cc3c(cc2F)C(=O)C(=C4SC(C)N34)C(=O)O |
| 9 | Prulifloxacin | CC1SC/2=C(C(O)=O)/C(=O)C3=CC(=C(C=C3N12)N4CCN(CC4)C\C5=C(C)\OC(=O)O5)F |
| 10 | NH125 | CCCCCCCCCCCCCCCC[N+]1=C(C)[N](CC2=CC=CC=C2)C=C1 |
| 11 | PicolinicAcid | O=C(O)c1ccccn1 |
| 12 | AgaricAcid | O=C(O)C(CCCCCCCCCCCCCCCC)C(O)(CC(=O)O)C(=O)O |
| 13 | Toltrazuril(sulfone) | CN1C(=O)NC(=O)N(C1=O)C2=CC=C(OC3=CC=C(C=C3)[S+]([O-])(=O)C(F)(F)F)C(=C2)C |
| 14 | 9(Z)-HexadecenoicAcid | CCCCCC\C=C/CCCCCCCC(O)=O |
| 15 | BenzylhexadecyldimethylammoniumChloride | [Cl-].C[N+](C)(Cc1ccccc1)CCCCCCCCCCCCCCCC |
| 16 | Corilagin | OC1C2COC(=O)C3=CC(=C(O)C(=C3C4=C(O)C(=C(O)C=C4C(=O)OC1C(O)C(O2)OC(=O)C5=CC(=C(O)C(=C5)O)O)O)O)O |
| 17 | Abametapir | CC1=CN=C(C=C1)C2=NC=C(C)C=C2 |
| 18 | Bictegravir | OC1=C2N(CC3OC4CCC(C4)N3C2=O)C=C(C(=O)NCC5=C(F)C=C(F)C=C5F)C1=O |
| 19 | Mebendazole | O=C(OC)Nc1nc2cc(ccc2n1)C(=O)c3ccccc3 |
| 20 | Fenbendazole | O=C(OC)Nc1nc2cc(ccc2n1)Sc3ccccc3 |
| 21 | EnalaprilMaleate | OC(=O)[C@@H]2CCCN2C(=O)[C@H](C)N[C@@H](CCc1ccccc1)C(=O)OCC |
| 22 | Entacapone | Oc1cc(\C=C(/C#N)C(=O)N(CC)CC)cc([N+]([O-])=O)c1O |
| 23 | Dichloro(ethylenediamine)platinum(II) | [Cl-][Pt]1([Cl-])NCCN1 |
| 24 | ElaidicAcid | CCCCCCCC/C=C/CCCCCCCC(O)=O |
| 25 | Dolutegravir(GSK1349572) | C[C@@H]1CCO[C@H]2CN\3/C=C(/C(=O)NCC4=C(F)C=C(F)C=C4)C(=O)\C(=C3C(=O)N12)O |
| 26 | Dolutegravir(GSK1349572) | C[C@@H]1CCO[C@H]2CN\3/C=C(/C(=O)NCC4=C(F)C=C(F)C=C4)C(=O)\C(=C3C(=O)N12)O |
| 27 | Cabotegravir | C[C@H]1CO[C@@H]2CN\3/C=C(/C(=O)NCC4=C(F)C=C(F)C=C4)C(=O)\C(=C3C(=O)N12)O |
| 28 | Cangrelor(AR-C69931) | CSCCNC1=NC(=NC2=C1N=C[N]2[C@@H]3O[C@H](CO[P](O)(=O)O[P](O)(=O)C(Cl)(Cl)[P](O)(O)=O)[C@@H](O)[C@H]3O)SCCC(F)(F)F |
